# Intraspecific diversity loss in a predator species alters prey community structure and ecosystem functions

**DOI:** 10.1101/2020.06.10.144337

**Authors:** Allan Raffard, Julien Cucherousset, José M. Montoya, Murielle Richard, Samson Acoca-Pidolle, Camille Poésy, Alexandre Garreau, Frédéric Santoul, Simon Blanchet

## Abstract

Loss in intraspecific diversity can alter ecosystem functions, but the underlying mechanisms are still elusive, and intraspecific biodiversity-ecosystem function relationships (iBEF) have been restrained to primary producers. Here, we manipulated genetic and functional richness of a fish consumer (*Phoxinus phoxinus*), to test whether iBEF relationships exist in consumer species, and whether they are more likely sustained by genetic or functional richness. We found that both genotypic and functional richness affected ecosystem functioning, either independently or in interaction. Loss in genotypic richness reduced benthic invertebrate diversity consistently across functional richness treatments, whereas it reduced zooplankton diversity only when functional richness was high. Finally, both losses in genotypic and functional richness altered essential functions (e.g. decomposition) through trophic cascades. We concluded that iBEF relationships lead to substantial top-down effects on entire food chains. The loss of genotypic richness impacted ecological properties as much as the loss of functional richness, probably because it sustains “cryptic” functional diversity.

## Introduction

Human disturbances associated with global change are increasingly altering worldwide patterns of species diversity, as well the functions and services provided by ecosystems (Parmesan 2006; Naeem *et al.* 2009; Isbell *et al.* 2017). Nonetheless changes observed at the species and ecosystem levels are always preceded by changes in phenotypic and genotypic composition within plant and animal populations (Clements & Ozgul 2016; Clements *et al.* 2017; Baruah *et al.* 2019). Accordingly, extremely rapid changes in intraspecific diversity are currently occurring worldwide (Darimont *et al.* 2009; Mimura *et al.* 2016; Miraldo *et al.* 2016; Leigh *et al.* 2019). Changes in intraspecific diversity can affect species turnover and composition (Vellend 2005; Vellend & Geber 2005), as well as ecosystem functioning (Des Roches *et al.* 2018; Raffard *et al.* 2019c). For instance, the loss of genotypes within primary producers can reduce ecosystem process rates and species diversity (Crutsinger *et al.* 2006; Johnson *et al.* 2006; Fridley & Grime 2010), suggesting the existence of biodiversity-ecosystem functioning relationships at the intraspecific level (iBEF, Hughes *et al.* 2008; Koricheva & Hayes 2018; Raffard *et al.* 2019c), in addition to the more widely studied BEFs at the interspecific level.

The relationships between intraspecific diversity and ecosystem functioning are challenging to study and this is primarily because of the confusion between genetic and functional trait diversity. iBEF studies have initially manipulated genotypic richness within experimental populations (e.g. Crutsinger *et al.* 2006; Hughes & Stachowicz 2009; Abbott *et al.* 2017). Genetic variability is expected to encapsulate a large proportion of trait variability, and thus higher genotypic richness should maintain higher functional diversity (Hughes et al. 2008). Although this approach allows deciphering the general effects of intraspecific diversity on ecosystem functioning, it does not provide mechanisms since ecological interactions are supported by functional traits that are -partly-genetically encoded. As a consequence, parallel studies have directly manipulated functional trait richness within populations, which enabled to determine a direct mechanistic linkage between functional richness and community structure and ecosystem functioning (Violle *et al.* 2007; Matthews *et al.* 2011). For instance, individual body mass (and traits covarying with body mass) has strong ecological effects because of the associated functional differences and resource use complementarity among individuals (Woodward *et al.* 2005; Hildrew *et al.* 2007; Rudolf *et al.* 2014; Harmon *et al.* 2009; Rudolf & Rasmussen 2013a). Focusing on specific traits, such as body mass, might conversely blur the ecological effects of “cryptic” trait variation (i.e. unmeasured functional traits such as metabolic rate or feeding behaviour and activity, Brown *et al.* 2004; Wolf & Weissing 2012) that is likely supported by genotypic richness. We therefore argue that manipulating simultaneously genotypic richness and the diversity of key functional traits such as body mass should allow assessing whether cryptic trait diversity is ecologically important, and could provide a better mechanistic understanding of iBEF relationships.

The ecological effects of biodiversity changes can be particularly strong when the later occur at high trophic levels such as secondary consumers or predators (Duffy 2002, 2003; Griffin *et al.* 2013). Changes in the diversity and abundance of predatory species can trigger important effects in functions supported by lower trophic levels, especially in regulating the abundance of the lower trophic levels and indirectly ecosystem functioning, such as biomass production (Shurin *et al.* 2002; Barnes *et al.* 2018). High predator species richness can sometimes favour resource use complementarity and decrease prey abundance (Cardinale *et al.* 2006; Griffin *et al.* 2013). In some cases, however, increasing predator richness increases prey abundance through mechanisms such as predator interference (Sih *et al.* 1998; O’Connor & Bruno 2009), and can modify multiple ecosystem functions along food webs (Antiqueira *et al.* 2018; Barnes *et al.* 2018). These mechanisms have all been investigated at the interspecific level, but they may also apply *within* a predatory species (intraspecific diversity), and we can expect relationships –either positive or negative-between predator intraspecific diversity and the structure of prey communities, that could subsequently cascade down on ecosystem functions such as decomposition rate or primary production (Duffy, 2002; Bruno & O’Connor, 2005). However, it is still difficult to forecast how loss in intraspecific diversity in consumer species could affect ecosystem functioning, because iBEF studies have primarily focused on primary producers (Koricheva & Hayes 2018; Raffard *et al.* 2019c). This is despite the fact that human activities strongly affect predator and consumer populations, for instance through harvest or fisheries activities, which may alter ecosystem functioning through intraspecific changes in traits and genotypes (Hauser *et al.* 2002; Estes *et al.* 2011; Palkovacs *et al.* 2012, 2018).

In this study, we investigated whether a loss in genotypic and functional diversity within a consumer species at the top of a three-level trophic chain could mediate top-down effects on key ecosystem functions. In a nine-months pond mesocosm experiment, we simultaneously manipulated *genotypic richness* (number of genetic entities) and the *functional richness* (variance in individuals body mass) of experimental populations of a freshwater fish, the European minnow (*Phoxinus phoxinus*); a common and abundant species with important ecosystems effects (Miró *et al.* 2018; Raffard *et al.* 2019b). We predicted that increased functional richness should affect ecological functions; if functional richness captures the entire functional differences among genotypes, then increasing genotypic richness should not impact ecological functions further (Fig. 1a). Alternatively, if functional richness does not capture all the functional differences among genotypes, then increasing genotypic richness should increase functional diversity and affect ecological functions (Fig. 1b). The ecological effects of genotypic and functional richness might display different shape following different conceptual predictions to which we compared our experimental findings (Fig. 1): (i) “additive” effects when loss in genotypic richness affects ecological functions regardless of changes in functional richness, (ii) “enhancing” effects when loss in genotypic richness affect ecological functions solely at higher levels of functional richness, or (iii) “compensatory” effects when high genotypic richness compensates for the loss of functional richness, maintaining higher ecological functions at lower levels of functional richness. Finally, we investigated the mechanistic basis of the ecological effects of genotypic and functional richness. In particular, we expected that genotypic and functional richness affect directly community structure through trophic interactions, and indirectly ecosystem functions at lower trophic levels through trophic cascades. Hence, we specifically tested whether the loss in genotypic and functional richness directly affected population performance and community structure, and indirectly ecosystem functioning mediated by changes in community structure of benthic and pelagic food web.

**Fig. 1.**
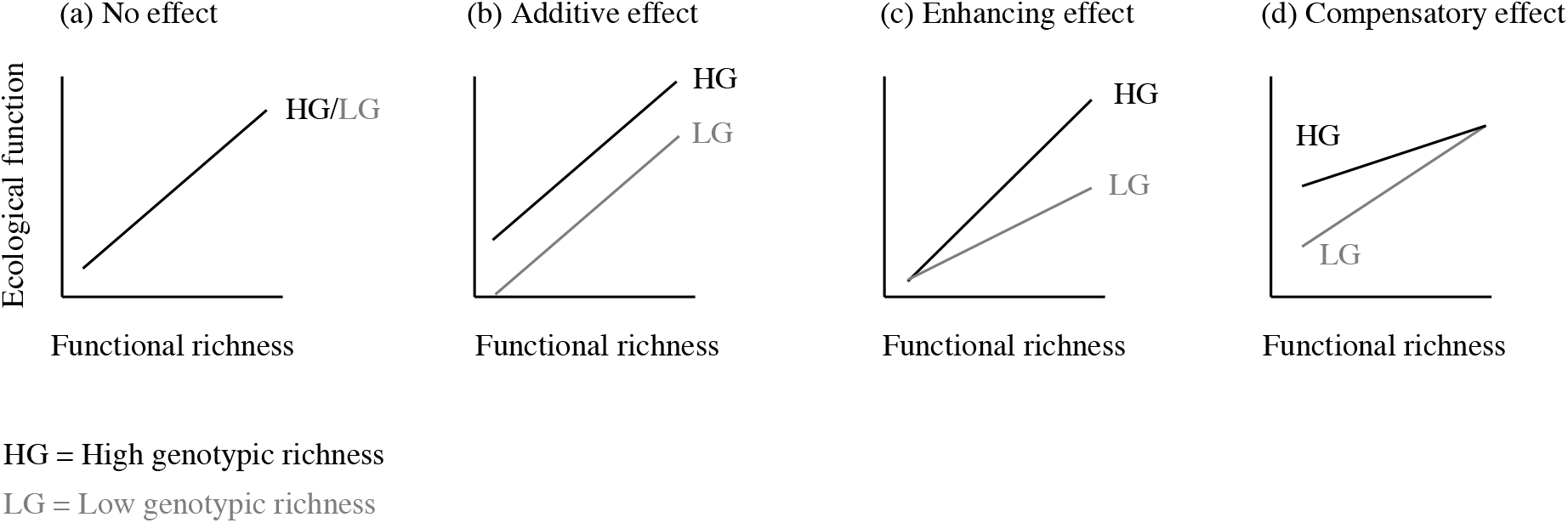
Predicted ecological effects of genotypic richness in relation with functional richness. Functional richness is assumed to positively affect ecological functions, whereas genotypic richness might have different effects depending on it support cryptic functional diversity. First, if functional richness does capture the essential of the functional differences among genotypes, then genotypic richness would have no ecological effects (a). Second, if genotypic richness supports cryptic functional diversity, the effects could be (b) additive, (c) enhancing, or (d) compensatory.

## Methods

### Model species

The European minnow (*P. phoxinus*) is a common species occurring across Western Europe, living in relatively cold water (summer water temperature generally lower than 24°C) including mountains lakes, small rivers at intermediate altitude and mountain streams (Keith *et al.* 2011). It is a small-bodied cyprinid fish species (<12 cm long, 4-8 cm long as an adult in general) playing the role of secondary consumer in most ecosystems (Raffard *et al.* 2020), with a generalist diet composed of small invertebrates, algae or zooplankton (Frost 1943; Collin & Fumagalli 2011). Populations of European minnows differed in their genetic and phenotypic richness (Fourtune *et al.* 2018), and previous works revealed that genetically and phenotypically unique populations differently affect prey community abundance and ecosystem functions (Raffard *et al.* 2019b).

We selected ten populations from a large river basin in southern France (the Garonne catchment) based on *a priori* knowledge to maximise both genetic and functional differentiations among populations (Fourtune *et al.* 2018; Raffard *et al.* 2019a) (Fig. S1). Specifically, the ten selected populations displayed a high level of genetic differentiation (mean pairwise Fst = 0.133, range = 0.029-0.320) and greatly varied in the number of alleles they harbour (from 5.470 alleles in average for the less diverse population to 10.176 alleles for the most diverse population) according to data measured using seventeen microsatellite markers (from Fourtune *et al.* 2018; Prunier *et al.* 2019; Raffard *et al.* 2019a). Moreover, we selected five populations mainly composed of small-bodied adults (mean body mass ± standard error (SE) = 1.03 g ± 0.02) and five populations mainly composed of large-bodied adults (mean body mass ± SE = 3.06 g ± 0.07) (Fig. S2a). These differences in adult body mass are due to selective pressures from the local environment such as mean water temperature; the higher the mean water temperature, the smallest the adult body mass due to increased metabolic rate and an accelerated pace-of-life (Raffard *et al.* 2019a). Finally, populations are geographically distant one from each other, and they will hereafter be each considered as functionally and genetically unique entity, and intraspecific richness will be manipulated by varying the number of populations in independent mesocosms (see details below).

In September 2017, individuals were collected in each river by electrofishing a 200-m section and visually selecting 30-50 individuals reflecting the size range of adults. Fish were maintained at similar density in outdoor tanks for 3 weeks before the start of the experiment. Fish were fed ad libitum with frozen Chironomidae during this period.

### Mesocosm experiment

In October 2016, 24 outdoor mesocosms were filled with 900 L of water, and 3 cm of gravel. Nutrients were added to the mesocosms using 10 mL of solution containing nitrogen and phosphorus (ratio N:P:K = 3.3:1.1:5.8). Each mesocosm was then inoculated twice (October 2016 and May 2017) with 200 mL of a concentrated solution of phytoplankton and 200 mL of concentrated solution of zooplankton collected from a unique lake nearby (Lamartine Lake, France 43°30′21.5″N, 1°20′32.7″E). In May 2017, we introduced three adult pond-snails (Lymnaeidae) and ten adult isopods (Asellidae) in each mesocosms. They were let uncovered to allow natural colonization by other invertebrates and community assemblage until the start of the experiment that occurred approximately 6 months later.

In October 2017, eight fish were weighted and introduced in each mesocosm, which were assigned to one of four treatments according to a full-factorial design with *genotypic* richness (two levels, *high* and *low* genotypic richness) and *functional* richness (two levels, *high* and *low* functional richness) as the main factors. Genotypic richness was manipulated by introducing individuals sourced from either two (*low genotypic richness*, four individuals from each population) or four (*high genotypic richness*, two individuals from each population) out of the ten populations displaying significant genetic differentiation (Fourtune *et al.* 2018; Raffard *et al.* 2019a). These two levels of genotypic richness were selected as it has previously been shown in a meta-analysis that the effect of intraspecific richness on ecosystem functioning increases linearly until approximately four genotypes and then reaches a plateau beyond that (Raffard *et al.* 2019c). Since we aimed at testing the effect of richness while minimizing the effects of population identity, each replicate of each treatment of richness contained a different (unique) assemblage of populations. Functional richness consisted in manipulating the body mass of the individuals present in the mesocosms; hence, experimental populations contained individuals sourced either from large or small populations (i.e., one functional entity, *low functional richness*), or from both small and large populations (i.e., two functional entities, *high functional richness*, see Table S1 and Fig. S2 for details on the different experimental populations). It is noteworthy that populations containing only either small or large individuals were actually more functionally redundant than populations containing large and small individuals. Experimental populations hence varied simultaneously according to their genotypic and functional richness, leading to four treatments.

The experiment lasted 30 weeks (210 days) after fish were introduced. Mesocosms were checked daily for mortality, which was rarely observed over the course of the experiment. At the end of the experiment, we measured several ecological parameters to assess effects of genotypic and functional richness on population performance, community structure and ecosystem functioning:

#### Population performance

All tanks were emptied and we recaptured all remaining fish to estimate mortality rate (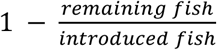 ; mean per tank ± standard error (SE) = 0.22 ± 0.01). Fish were weighted to assess fish biomass production (biomass production = averaged final weight - averaged initial weight) of each experimental population during the experiment. Since individuals with small initial body mass intrinsically displayed a higher growth rate than large individuals, a standardised biomass production index was calculated as the residuals of the relationship between biomass production and initial fish size.

#### Community structure

Zooplankton community structure was quantified by filtering 5 L of water through a 200 μm sieve. Samples were conserved in a 70% ethanol solution and subsequently identified to the order or family levels, including Cyclopoida, Calanoida, Daphniidae, Chydoridae and Bosminidae. The diversity of zooplankton was calculated as the Simpson’s diversity (*D-zoo*) representing the probability that two randomly chosen individuals belong to different clades. *D-zoo* was calculated as 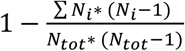, where *N*_*tot*_ was the total number of sampled individuals, and *N*_*i*_ the number of sampled individuals for each group (Simpson 1949; Lande 1996). Zooplankton abundance was quantified as the total number of individuals for all taxa pooled at the tank level.

Benthic invertebrates were collected from the mesh bags used to measure decomposition rates (*see below*), conserved in a 70% ethanol solution, and identified as Isopoda, Diptera, Gastropoda, Ephemeroptera, Plecoptera, Odonata, Copepoda, Cladocera, and Ostracoda. The diversity of benthic invertebrates was calculated as the Simpson’s diversity (*D-inv*). The abundance of benthic invertebrates was quantified as the total number of individuals for all taxa pooled at the tank level.

#### Ecosystem functioning

Decomposition rate was measured by quantifying the mass loss of black poplar (*Populus nigra*, a dominant riparian tree in southern France) abscised leaves (Alp *et al.* 2016). On the 7^th^ of March 2018 (19 weeks after the introduction of fish), 4 g of air-dried leaves were put in each mesocosm within a coarse mesh (1 × 1 cm) bag. At the end of the experiment, the remaining leaf material was removed from the mesocosms, rinsed with tap water, oven dried at 60°C for three days and weighed to the nearest 0.001 g to assess the loss of biomass. The decomposition rate was calculated as 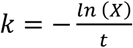 (Alp *et al.* 2016), where *X* is the proportion of litter remaining and *t* is the elapsed time in days.

Pelagic algae stock was measured as the chlorophyll-a concentration (μg.L^−1^) in the water column using a multiparametric probe (OTT, Hydrolab DS5^®^). Five measurements were taken in each mesocosm and averaged for subsequent analyses. Since phytoplankton are the main primary producers in these mesocosms (Downing & Leibold 2002), pelagic algae stock can be considered as a proxy for the biomass of primary producers.

### Statistical analyses

Prior to analysis, the pelagic algae stock (i.e. the chlorophyll-a concentration), zooplankton and benthic invertebrate abundances were log-transformed to reach normality of models’ residuals. After analyses of outliers using Cook’s distance, we removed one mesocosm from the final analyses that displayed outliers for several of the measured variables (Fig. S3). Analyses were hence run on 23 replicates.

We first assessed the ecological effects of genotypic and functional richness using mixed effects linear models (LMMs). We ran one model for each ecological parameter as a dependent variable, and the genotypic richness, functional richness, and the resulting two-term interaction (that was removed when non-significant) as explanatory variables. Fish mortality rate during the experiment was included as a fixed effect to control for a potential effect of final density on ecological processes. To control for the disposition of the tanks during the experiment, the position of tanks was added as a random term. To compare the relative strength of a loss in genotypic and functional richness on ecological parameters we calculated effect sizes. Hedges’ g were computed as follow: 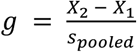, where *X*_*1*_ and *X*_*2*_ are means of treatments (for each genotypic or functional richness treatment separately) measured for each ecological parameters respectively, and s_pooled_ is the pooled standard deviation computed as 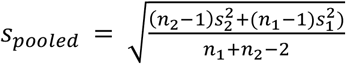, where n is sample size, and s^2^ is variance (Nakagawa & Cuthill 2007). An effect size was hence measured for each variable response (n = 7, and each treatment) independently. These individual effect sizes were then averaged for genotypic (*g*_*G*_) and functional richness (*g*_*S*_) respectively so as to get an absolute mean effect sizes that were compared visually based on 95% confidence intervals (CI).

Secondly, we ran confirmatory path-analysis (Shipley 2009) to assess the direct and indirect links among intraspecific diversity (genotypic and functional richness), prey (invertebrates and zooplankton) community structure and ecosystem functioning. We expected that intraspecific diversity in European minnows affects directly invertebrate community (both diversity and abundance) through predation, and indirectly ecosystem functions (decomposition rate, algae stock) through trophic cascades. Specifically, benthic invertebrates consume leaf litter, affecting decomposition rate (Gessner *et al.* 1999, 2010), while zooplankton forages on phytoplankton, affecting pelagic algae stock (brown and green trophic food chain, respectively). Because genotypic and functional richness likely interact to shape processes (see Fig. 1), we initially included in all models the interaction term between genotypic and functional richness on community structure. As above, interaction terms were removed when not significant. Specifically, we computed two path analysis: one for the benthic food chain (fish-invertebrates-decomposition), and one for the pelagic food chain (fish-zooplankton-algae stock) (Fig. S4). The fits of the structural models were assessed using the *C* statistic (see independent claims in Table S2), that follows a χ^2^ distribution and whose associated *p*-value indicates whether the models adequately fit the data or not. Alternative models including direct effects of fish intraspecific diversity on community components and ecosystem functions were performed (Fig. S4) to test whether genotypic and functional richness affected ecosystem functions through unmeasured parameters (e.g., other community parameters, nutrient recycling). This approach allows testing whether the predicted indirect effects are the most likely effects that explain the data. Finally, the structural models were further compared to simplified models that did not include the effects of genotypic and functional richness on community parameters (see Fig. S4) using Akaike Information Criteria (AIC, Cardon *et al.* 2011; Shipley 2013). This later test allows assessing whether the indirect effects of genotypic and functional richness lead to a better explanation of the data, and whether trophic cascades were important or not.

All statistical analysis were run using R software (R Core Team 2013), LMM were run using the R-package lme4 (Bates *et al.* 2014), and confirmatory path-analyses were run using the R-package ggm (Marchetti *et al.* 2020).

## Results

Both genotypic and functional richness of experimental populations significantly affected several ecological processes (Table 1). At the population level, we found that the interaction between genotypic and functional richness tended (p = 0.057) to alter fish biomass production (i.e., population performance) of experimental populations (Table 1, Fig. 2a). Specifically, the less diversified experimental populations (low genotypic richness and low functional richness) displayed a lower biomass production than all other treatments (Fig. 2a). At the community level, the diversity of benthic invertebrates was significantly higher in the high genotypic richness treatment (mean *D-inv* ± SE = 0.64 ± 0.04) than in the low genotypic richness treatment (*D-inv* ± SE = 0.53 ± 0.03), and this pattern was repeatable across functional richness treatments (Fig. 2d, Table 1). Irrespective of the genotypic richness treatment, mesocosms containing fish populations with low functional richness had higher diversity of benthic invertebrates than mesocosms containing fish populations with high functional diversity (Fig. 2d, Table 1). In contrast, the diversity of zooplankton (*D-zoo*) was affected by the interaction between genotypic and functional richness (Fig. 2e, Table 1). Zooplankton diversity was strikingly higher in populations with both high genotypic richness and high functional richness (Fig. 2e, Table 1), suggesting that, in this case, the effect of genotypic richness depended on the functional richness of experimental fish populations. Benthic invertebrates and zooplankton abundances were consistently (i.e. across genotypic richness treatment) enhanced when increasing functional richness (Fig. 2f, g, and Table 1). Genotypic richness of populations did not affect benthic invertebrates and zooplankton abundances (Fig. 2f, g, and Table 1). A weak but significant negative effect of functional richness on the pelagic algae stock was detected (Table 1, Fig. 2c) and there was no significant effects genotypic and functional richness on decomposition rate (Table 2). Overall, genotypic and functional richness induced ecological effects of similar intensity (mean |g_G_| = 0.356, CIs = 0.043-0.669, and mean |g_S_| = 0.564, CIs = 0.247-0.882, see Fig. S5 for details).

**Table 1.**
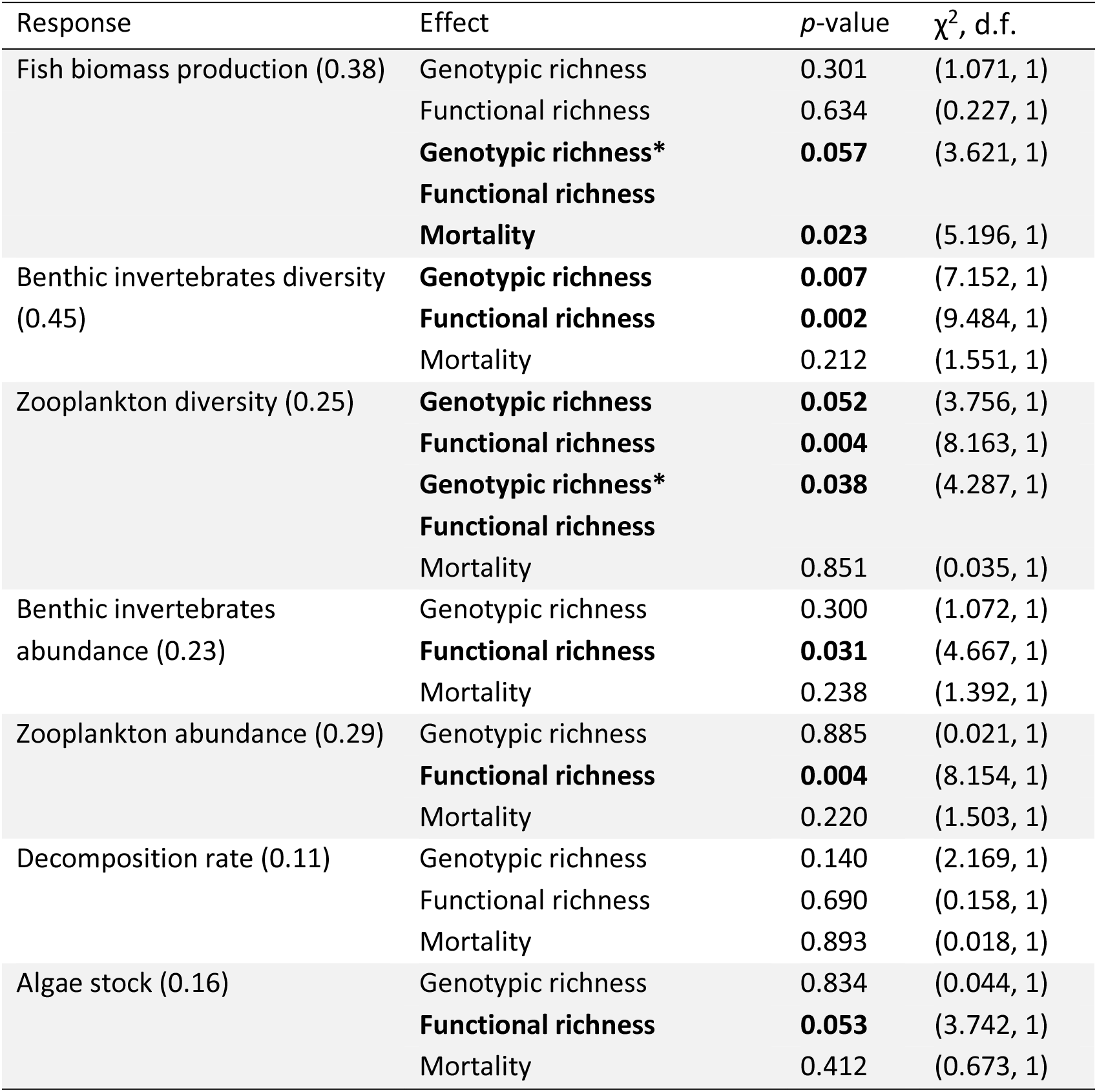
Results of the mixed effect linear models quantifying the relationships between genotypic richness, functional richness and ecological parameters. Significant -and near significant-*p*-values are displayed in bold, R^2^ are shown into brackets. Interaction terms were removed from the model when not significant

**Fig. 2.**
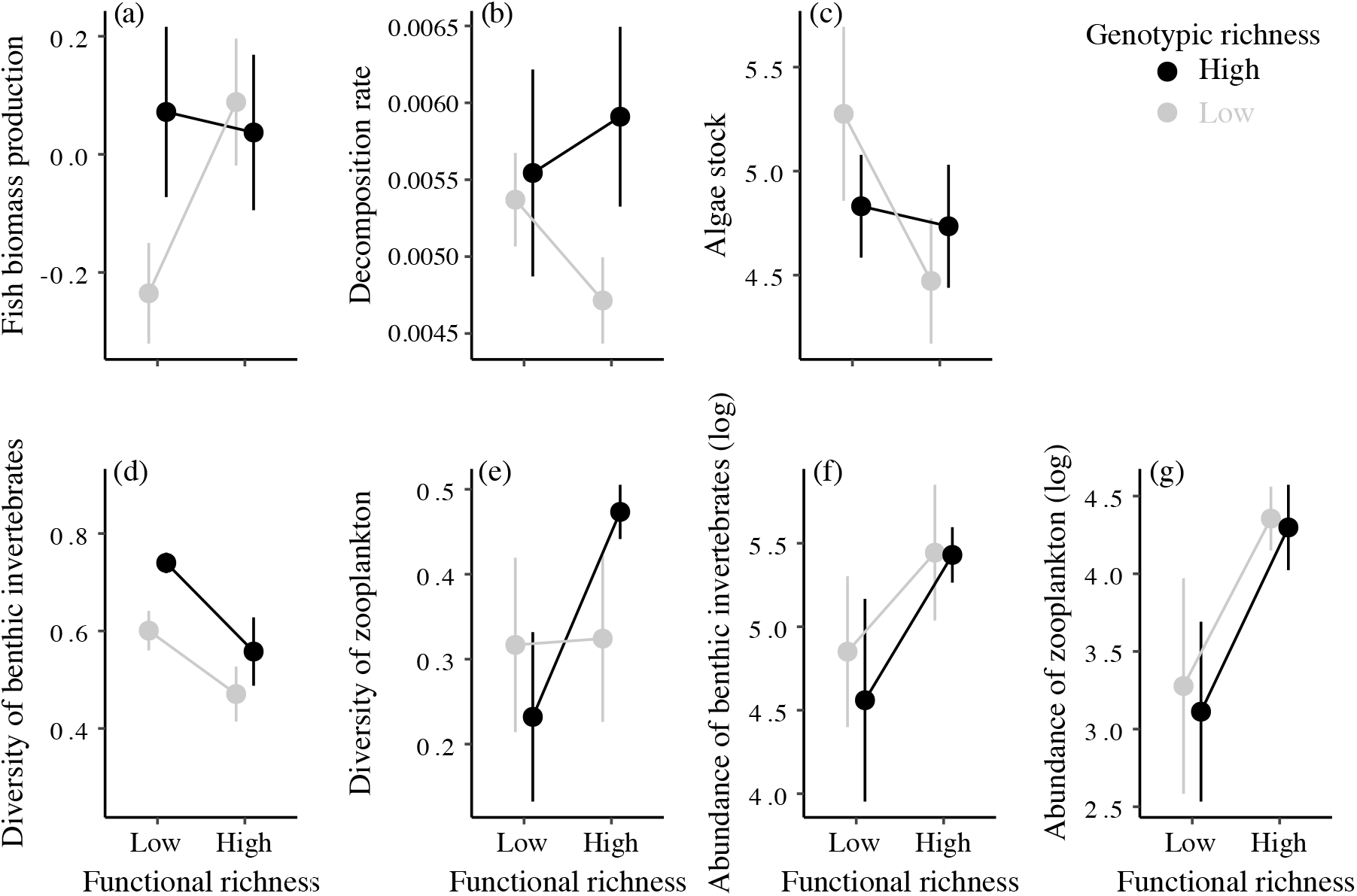
Effects of genotypic and functional richness on populations biomass production index (a), decomposition rate (b), algae stock (c), diversity of benthic invertebrates (e), diversity of zooplankton (f), abundance of benthic invertebrates (g), and abundance of zooplankton (h). Error bars represent ± 1 SE.

**Table 2.** Model fits of the two confirmatory path-analyses explaining decomposition rate and algae stock variation. C-statistic (*C*), degree of freedom (*df*), and *p*-value are given for full models as indication.

| Variable | $C$ | $df$ | $p$-value | AIC full model | AIC alternative model | AIC simplified model |
| --- | --- | --- | --- | --- | --- | --- |
| Decomposition rate | 8.555 | 6 | 0.200 | 26.555 | 30.555 | 53.042 |
| Algae stock | 6.745 | 10 | 0.749 | 26.746 | 28.367 | 49.679 |

Confirmatory path analyses revealed that effects of intraspecific diversity on ecosystem functioning were primarily mediated by trophic cascades, through changes in community structure (Fig. 3). First, genotypic and functional richness affected the diversity and abundance of benthic invertebrates (Fig. 3), which subsequently and positively affected the decomposition rate (*p* < 0.001, Fig. 4a, b and Table S3). Second, functional richness positively affected abundance of zooplankton, leading to a decrease in pelagic algae stock (*p* < 0.001, Fig. 4c, d, Table S3). Overall, the confirmatory path-analysis confirmed the key role of intraspecific diversity for both food chains, and its indirect effects for ecosystem functions (Fig. 3 and Table 2). Indeed, models including genotypic and functional richness reproduced adequately the causal pathways (Table 2), and their AICs were better than that of both alternative and simplified models for each ecosystem function (Table 2).

**Fig. 3.**
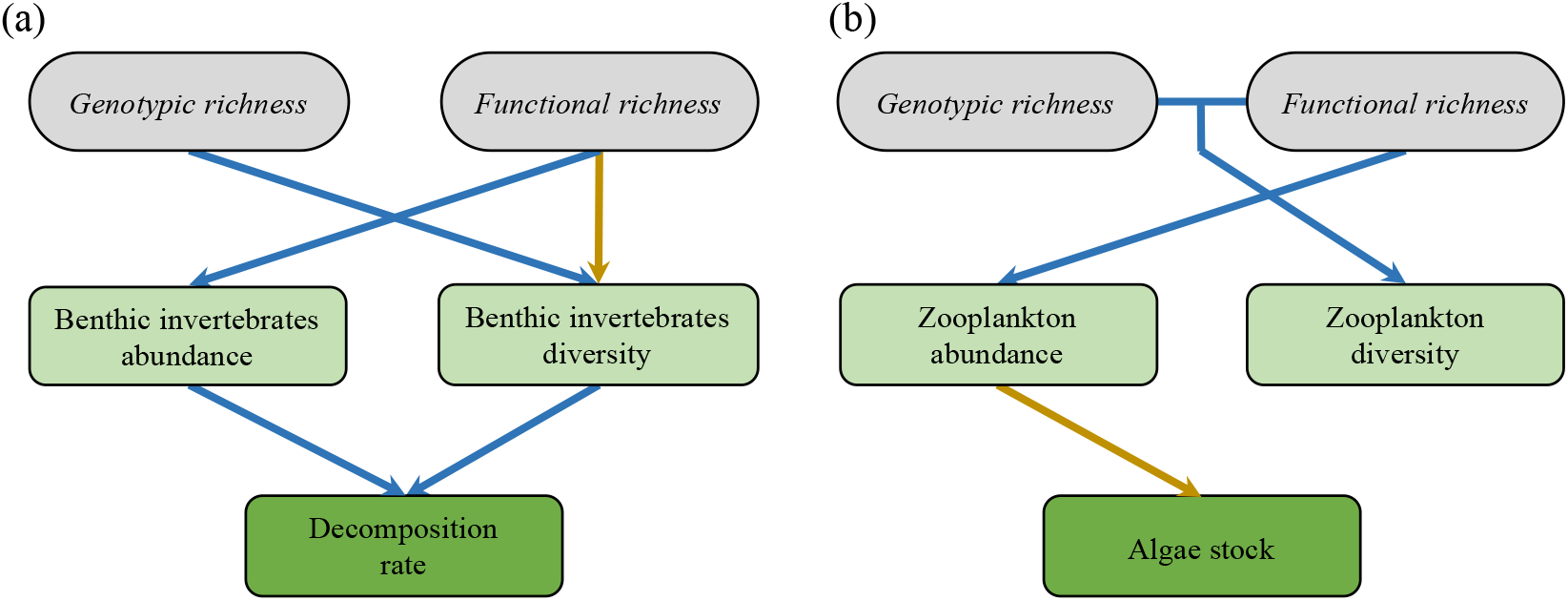
Causal pathways between genotypic richness, functional richness, community structure, and ecosystem functioning. Two models have been run separately on decomposition rate (a), and pelagic algae stock (b). Only significant links are drawn, blue arrows represent positive links, yellow arrows negative links.

**Fig. 4.**
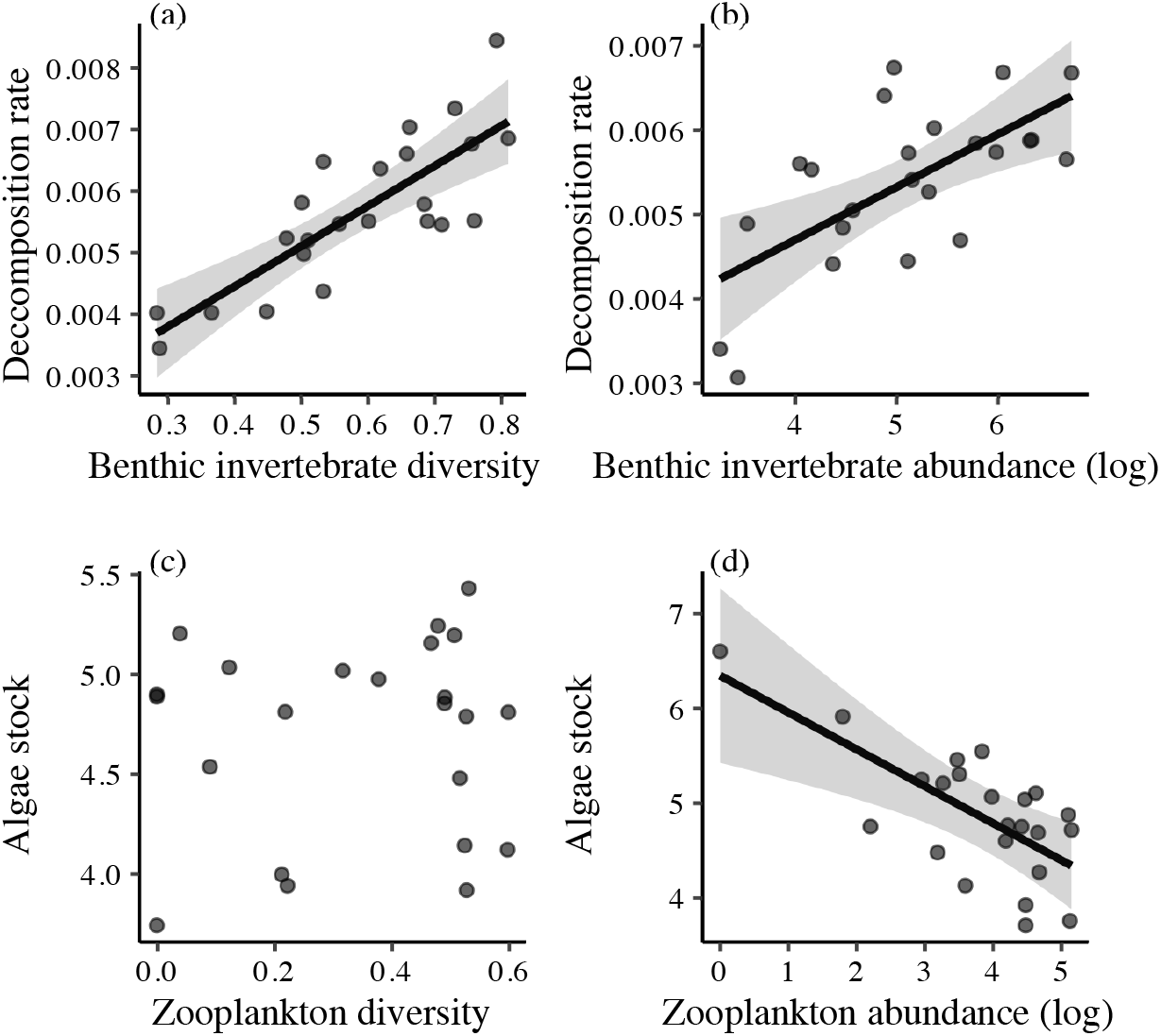
Effects of the diversity (a) and the abundance (b) of benthic invertebrates on decomposition rate; and effects of the diversity and the abundance of zooplankton on pelagic algae stocks (c and d, respectively). Points are partial residuals extracted from models described in the statistical analysis section (see also Table S3). Straight lines represent the slope predicted by the models (see statistical analyses) when significant (α < 0.05), and grey shadows represent 95% CIs.

## Discussion

The present study demonstrates that loosing intraspecific diversity in a secondary consumer species has substantial top-down consequences for community structure and ecosystem functioning in food webs. We showed that both losses in functional (mass) and genotypic richness sustained these iBEF relationships, suggesting that differences in mass among individuals (an important functional trait for food web dynamics, Woodward *et al.* 2005) did not capture the entire functional space, and thus that genotypic richness encapsulates important and “cryptic” (unmeasured) functional diversity. The loss of genotypes within consumer populations can affect both the community structure and the abundance of lower trophic levels, as well as ecosystem functions with a similar strength than the loss of functional entities (i.e. body mass). Specifically, we found that diversity loss (genotypic and functional) within populations indirectly affects primary producer biomass and organic matter recycling, two ecosystem functions forming the bases of food chains. This suggests that intraspecific diversity is a key but understudied facet of biodiversity as it indirectly sustains BEF relationships, even when changes in intraspecific diversity occur at the top of the food chains.

Our study suggests that genotypic richness can support non-negligible cryptic functional diversity. Cryptic functional traits, such as physiological and behavioural traits, can affect interactions in food webs and ecological functions. As such, behavioural variation within populations is widespread in nature, and can be genetically encoded (Sih *et al.* 2004; Anreiter & Sokolowski 2019). For instance, personality traits (e.g., activity or boldness) can determine foraging activity and diet, ultimately affecting trophic interactions and ecosystem functioning in food webs (Wolf & Weissing 2012; Toscano *et al.* 2016). Additionally, metabolic and stoichiometric traits, can also strongly differ within species, and can be functionally important in shaping energetic needs both quantitatively and qualitatively (i.e., specific elemental needs) (Brown *et al.* 2004; Leal, *et al.* 2016; Rosenblatt & Schmitz 2016). Hence, fish with different genotypes may share obvious functional traits (such as body mass) but may subtly differ in other cryptic functional traits, making them unique ecologically (Loreau 2001). Although such “ecological dissimilarity-despite-morphological similarity” has rarely been demonstrated within species, there is now ample evidence that cryptic species (species being morphologically and phylogenetically similar) can actually be ecologically unique regarding their influence on ecosystems (e.g. Fišer *et al.* 2015).

Interestingly, this cryptic diversity can interact with functional richness in various ways (i.e., additive, enhancing, or compensatory) depending on the ecological parameter considered (i.e., benthic invertebrate diversity, zooplankton diversity, and fish biomass production, respectively). For instance, we found that genotypic richness can compensate for the loss of functional diversity for population performance (i.e., increase fish biomass production), with high genotypic richness maintaining high biomass production even when the distribution of body mass in the population is limited. We speculate that genotypic richness can increase niche variation among individuals, thereby limiting the loss of biomass production when decreasing functional richness (Duffy *et al.* 2001). These findings echo and expand on studies at the interspecific level demonstrating that phylogenetic diversity among species explains variance in ecosystem functions that is not explained either by species or functional diversity, suggesting that phylogenetic diversity sustains unmeasured functional differences among species (Cadotte *et al.* 2012; Mouquet *et al.* 2012; Le Bagousse-Pinguet *et al.* 2019). We argue that iBEF relationships are sustained by differences in functional traits among individuals that can be directly measured (e.g. mass) and/or indirectly captured through quantification of the genetic pool of individuals composing populations.

Accordingly, we showed that genotypic and functional richness can affect independently (i.e., additively) and consistently community structure, demonstrating that multiple facets of intraspecific diversity can regulate lower trophic levels. First, genotypic richness increased benthic invertebrate diversity. This effect was very robust, as it was repeatable and consistent across functional richness treatments. Genotypic richness probably enhanced resource partitioning, allowing individuals to forage on a more diverse array of resources, regulating the abundance of each taxonomic group and leading to a higher diversity (Duffy, 2002; Johnson *et al.*, 2006; Hughes et al. 2008). Second, functional richness led to increased prey abundance that was repeatable across both communities of benthic invertebrates and zooplankton. Previous studies at the inter- and intraspecific levels provide non-consistent predictions, since an increase in consumer diversity can affect either positively or negatively resource abundance (Sih *et al.* 1998; Griffin *et al.* 2013; Rudolf & Rasmussen 2013; Antiqueira *et al.* 2018; Barnes *et al.* 2018). Our results indicate that functionally rich populations did not consume fewer resources than functionally poor populations since they had higher biomass production. This suggests that the increase in prey abundance was not due to an increase in inter-individual competition in functionally rich populations. At the opposite, flexible exploitation of resources might occur in functional rich populations because European minnows are omnivorous (Frost 1943; Collin & Fumagalli 2011), their diet probably included some periphyton, decreasing the predation risk on animal resources, and hence increasing invertebrate abundance. Because the top-down control of intraspecific diversity on community structure is likely driven by trophic mechanisms, quantifying individual diet (e.g. using gut content, eDNA or stable isotope analysis) and its temporal dynamic in such experiments would allow making more accurate predictions regarding trophic niche partitioning.

Our results further revealed how changes in top consumers’ genotypic and functional richness percolate through the food web and alter ecosystem functions at the base of the food chains. Such trophic cascade induced by intraspecific diversity could be driven by a “classical” (i.e. interspecific) BEF relationship between benthic invertebrate diversity and decomposition rate (Hooper *et al.* 2005; Gessner *et al.* 2010). Specifically, fish genotypic richness increased benthic invertebrate diversity that accelerated litter decomposition rate. The higher decomposition rate of organic matter is likely produced by higher consumption efficiency through trophic complementarity among clades of invertebrates in diverse community (Gessner *et al.* 2010). Invertebrate community with a high diversity probably harboured a high functional diversity (Cadotte *et al.* 2011), and focusing on the functional type of invertebrates might allow a more precise understanding of this link. These results echo those reported at the community level and those manipulating richness within primary producer species (Downing & Leibold 2002; Crutsinger *et al.* 2006), while implying a modification of top-down effects by intraspecific diversity in consumers. Biodiversity within consumer species is largely impacted by anthropogenic activities (Miraldo *et al.* 2016), involving the loss of non-measured functional diversity, which might further lead to underestimated cascading impacts on ecosystem functioning.

In conclusion, we demonstrated that both genetic and functional richness within consumer populations are important facets of biological diversity, inducing effects on prey community structure and trophic cascades mediating ecosystem functions. These results are consistent with previous synthetic works (Koricheva & Hayes 2018; Raffard *et al.* 2019c), reinforcing the call for considering changes of intraspecific diversity in consumer species as an important ecological predictor. Importantly, genotypic richness can sustain important cryptic functional diversity, and future investigations should aim at developing a general framework from genes to ecosystems to better understand the global links existing among the multiple facets of biodiversity and ecosystem functioning, and ultimately ecosystem services.

## Data accessibility

Data are available online: 10.6084/m9.figshare.12459065

## Supplementary material

Script and codes are available online: 10.6084/m9.figshare.12459065

## Acknowledgements

We thank Kéoni Saint-Pe and Jérôme G. Prunier for their help for fish sampling. AR is financially supported by a Doctoral scholarship from the Université Fédérale de Toulouse. Electrofishing was performed under the authorization of local authorities (Arrêtés Préfectoraux from the Direction Départementale des Territoires of French departments Ariège, Haute-Garonne, Tarn, Aveyron, Tarn-et-Garonne, Lot). This work was undertaken at SETE, which is part of the “Laboratoire d’Excellence” (LABEX) entitled TULIP (ANR-10-LABX-41). Version 3 of this preprint has been peer-reviewed and recommended by Peer Community In Ecology (https://doi.org/10.24072/pci.ecology.100064)

## Conflict of interest disclosure

The authors of this preprint declare that they have no financial conflict of interest with the content of this article. Simon Blanchet and José M. Montoya are one of the PCI Ecology recommenders.

## Appendix

**Fig. S1.**
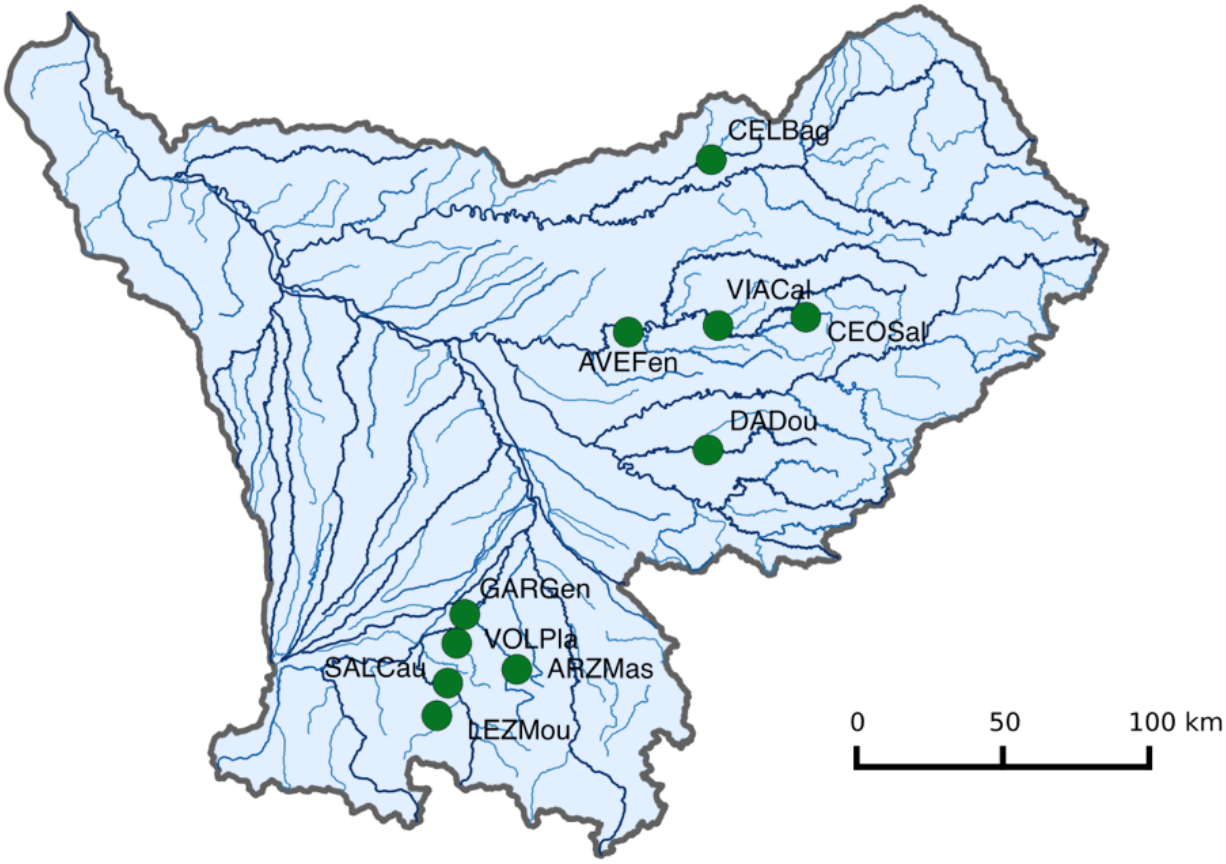
Geographical distribution of the ten populations of European minnows (*Phoxinus phoxinus*) used in this study.

**Fig. S2.**
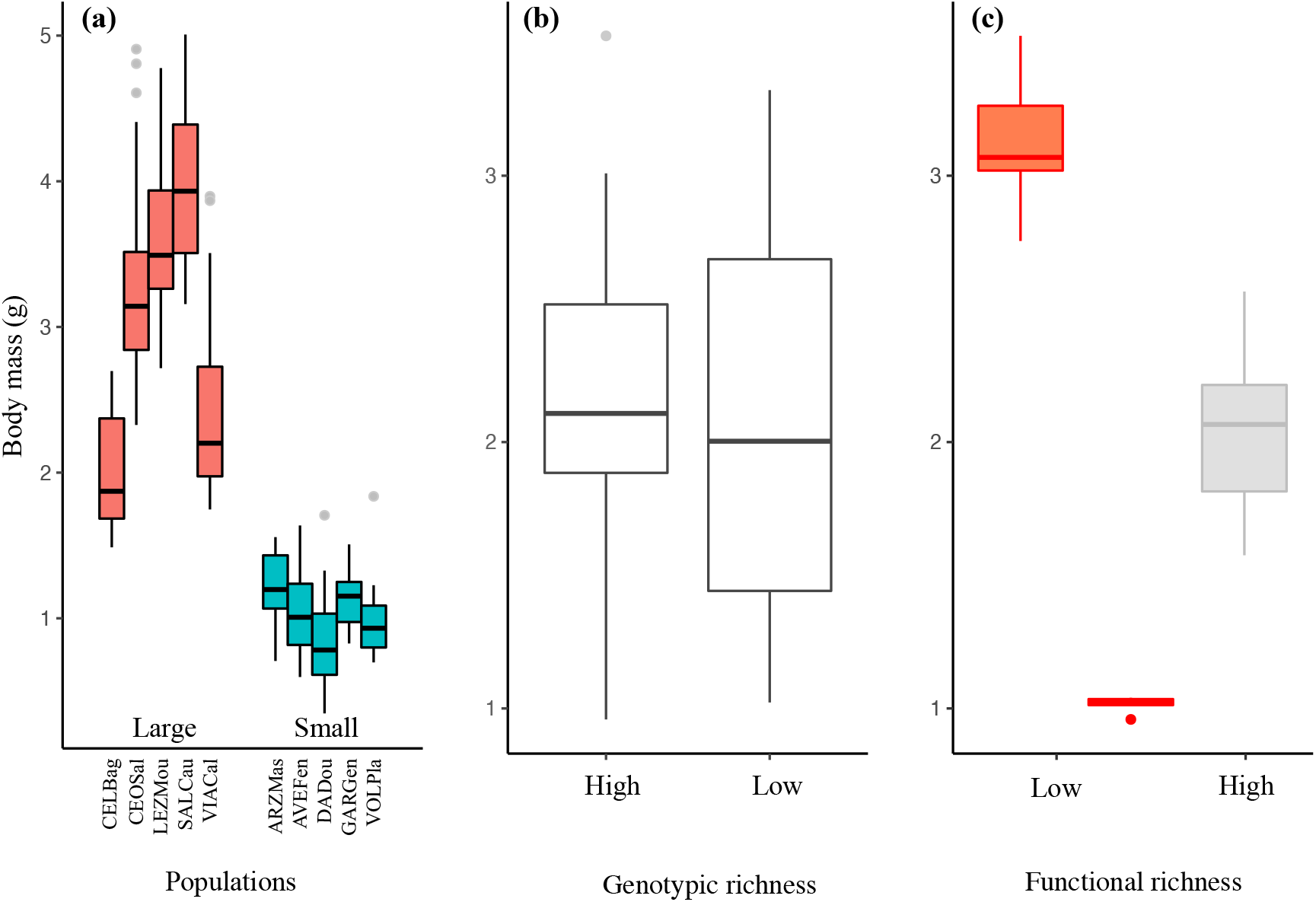
Relationships between fish body mass and **(a)** population of origin, **(b)** genotypic richness treatments (high = four populations, and low = two populations), and **(c)** functional richness.

**Fig. S3.**
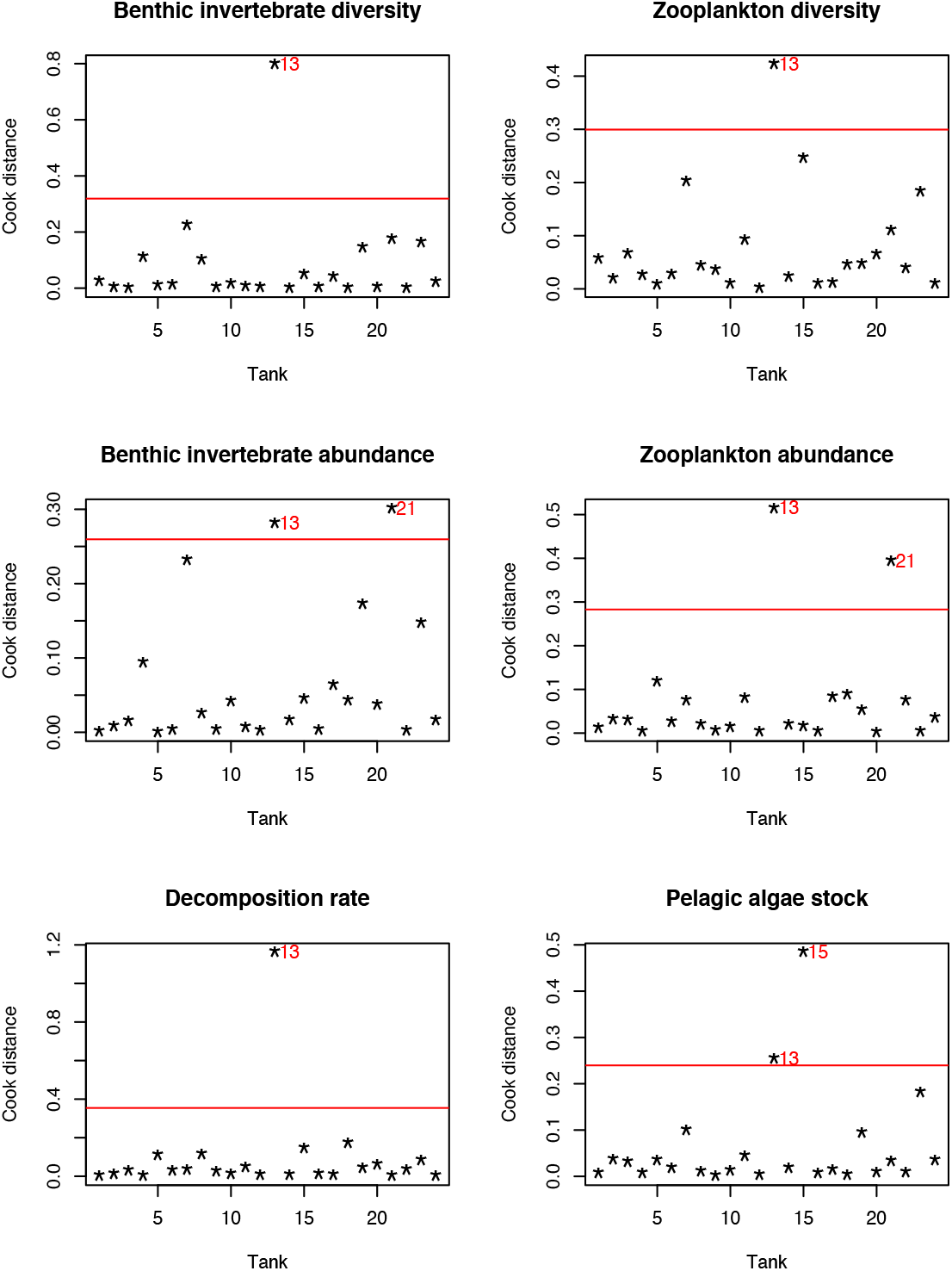
Analysis of outliers using Cook’s distance (Cook 1977; Cousineau & Chartier 2010). The higher the distance the more influential the points on the variable. The horizontal bar, representing the mean across all tanks multiplied by four, is given as an indicative threshold above which a point may be considered as influential. Tank 13 was influential in all variable, and was therefore discarded from analyses.

---

**Fig. S4.**
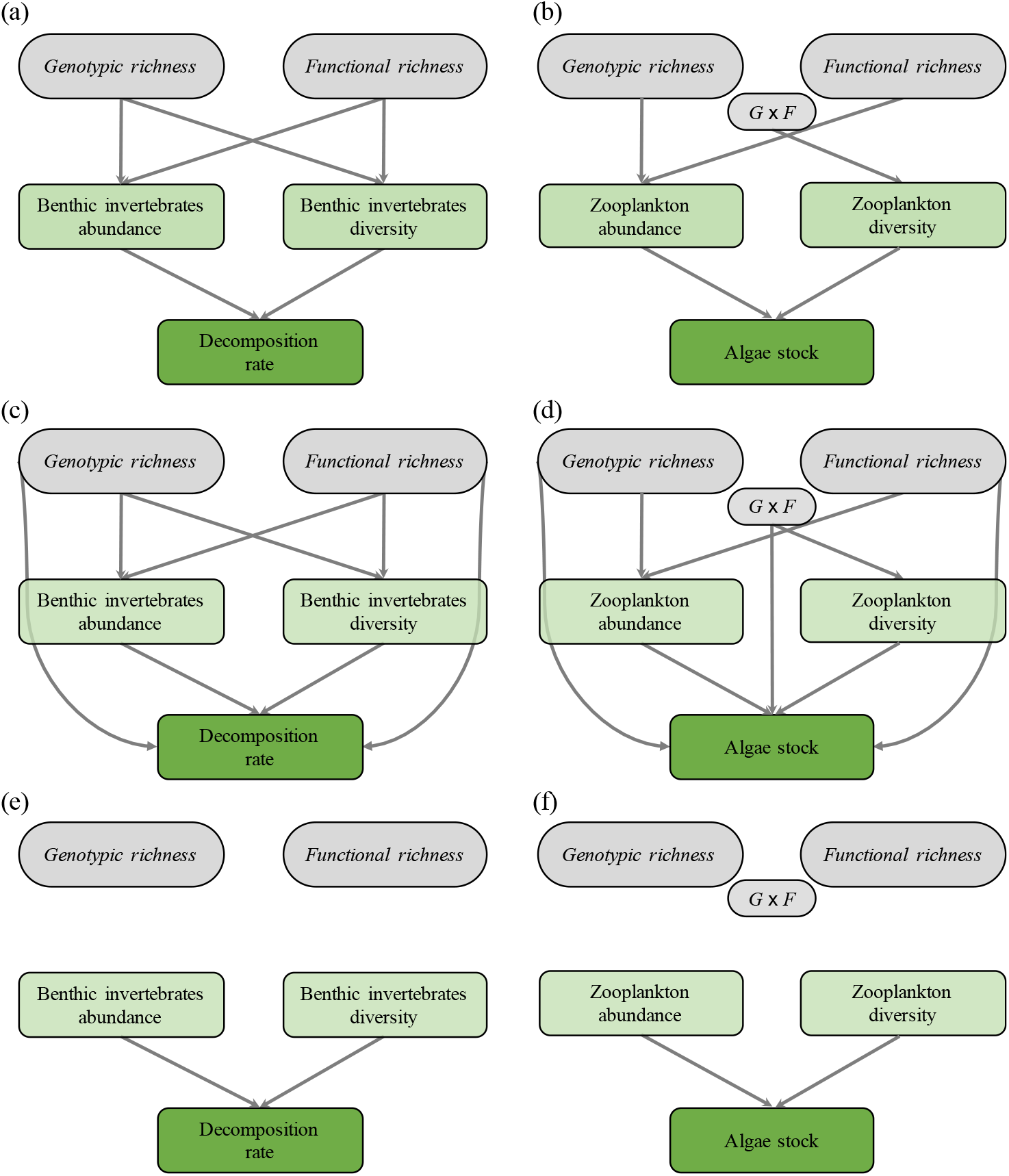
Diagram of the causal pathways used to explain variation in **(a)** decomposition rate and **(b)** algae stock. These models were compared to alternative models (**(c)** and **(d)**, respectively) including direct effect of genotypic and functional richness on ecosystem functions. Finally, simplified models were performed, in which the effects of genotypic and functional richness on community structure were excluded (**(e)** and **(f)**). ‘*G × F* ’ denotes the interaction between genotypic and functional richness.

**Fig. S5.**
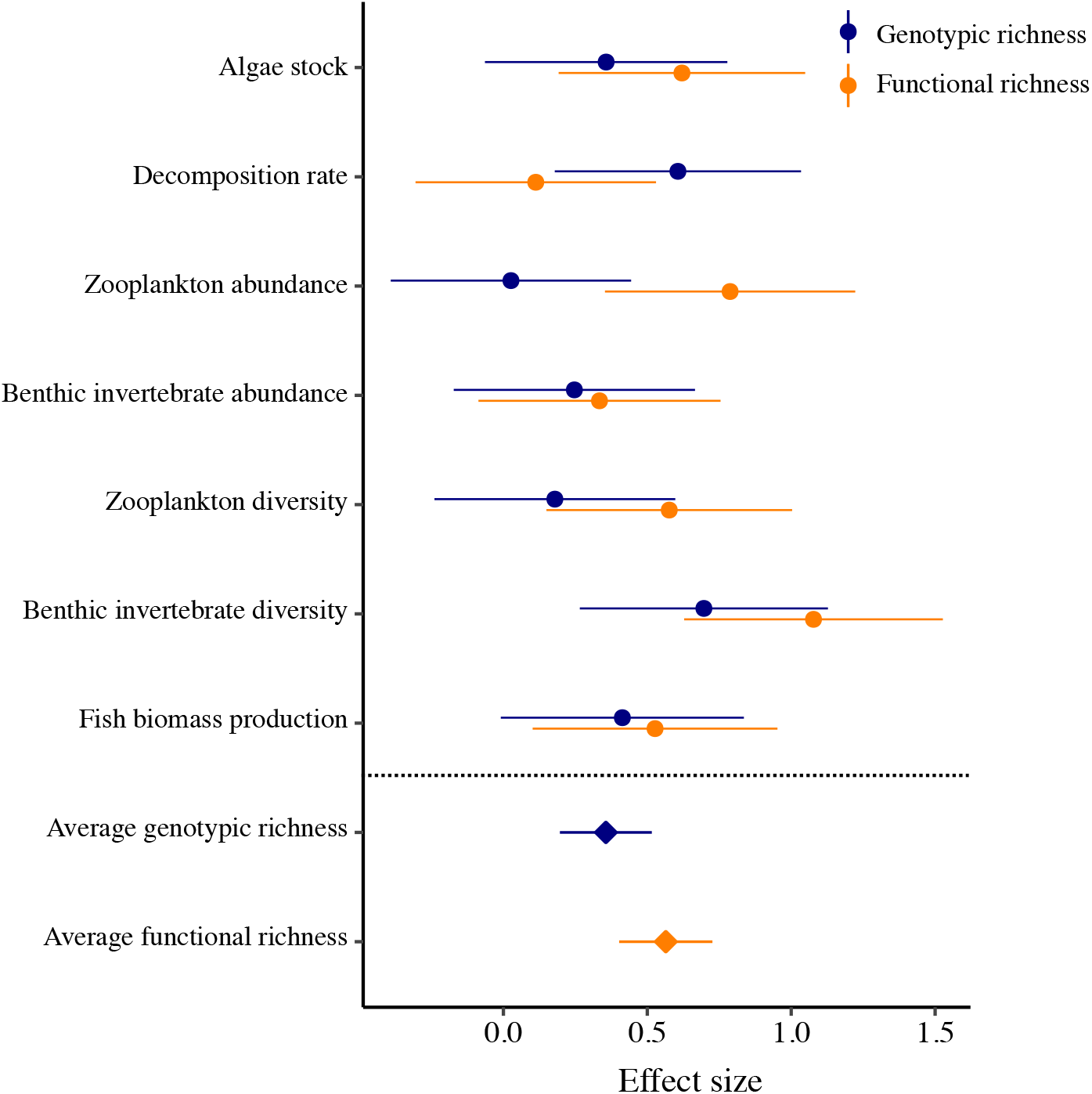
Effect size (Hedges’ g) of genotypic and functional richness for each ecological parameter and their overall means. Error bars represent ± 1SE.

**Table S1.** Population origin of minnows (*Phoxinus phoxinus*) used in each experimental treatment.

| Low genotypic richness<br>Low functional richness | Low genotypic richness<br>High functional richness | High genotypic richness<br>Low functional richness | High genotypic richness<br>High functional richness |
| --- | --- | --- | --- |
| CElBag-SALCau | CEOSal-VOLPla | CEOSal-VIASeg-CElBag-SALCau | CEOSal-CElBag-AVEfen-ARZMas |
| LEZMou-CEOSal | CElBag-GARGen | CEOSal-VIASeg-LEZMou-SALCau | CEOSal-VIASeg-AVEfen-VOLPla |
| VIACal-SALCau | VIASeg-ARZMas | CEOSal-VIASeg-LEZMou-CElBag | LEZMou-VIASeg-DADou-VOLPla |
| GARGen-AVEFen | CEOSal-AVEFen | GARGen-AVEFen-DADou-ARZMas | CElBag-VIASeg-ARZMas-VOLPla |
| VOLPla-ARZmas | LEZMou-DADou | VOLPla-AVEFen-DADou-ARZMas | LEZMou-SALCau-DADou-GARGen |
| DADou-AVEFen | SALCau-AVEfen | VOLPla-AVEFen-DADou-GARGen | SALCau-CEOSal-GARGen-AVEFen |

**Table S2.**
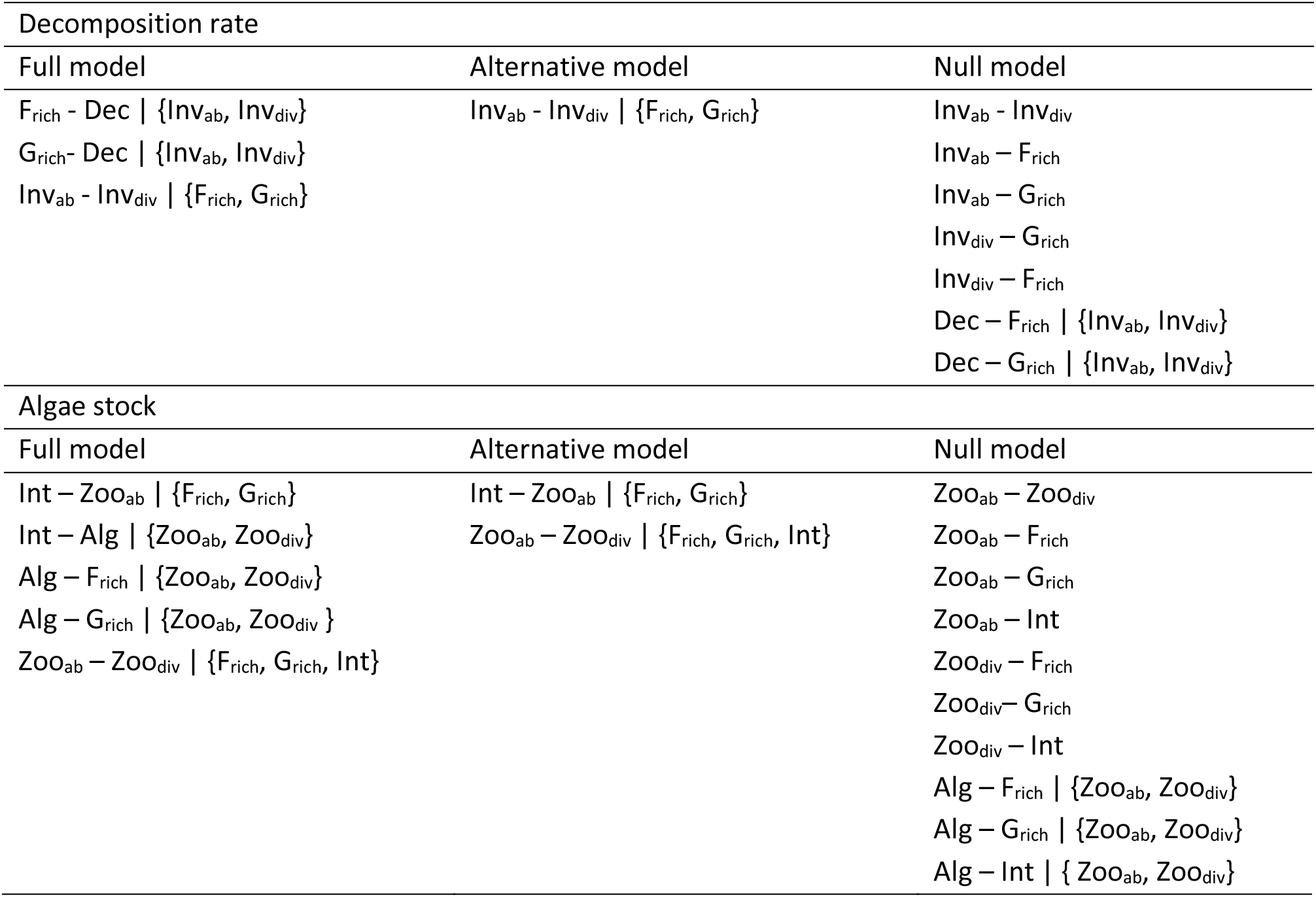
D-separation claims implied in the structural models shown in Fig. S4. Frich: functional richness; Grich: genotypic richness; Int: FxG interaction; Invab: benthic invertebrates abundance; Invdiv: benthic invertebrates diversity; Dec: decomposition rate; Zooab: zooplankton abundance; Zoodiv: zooplankton diversity; Alg: Algae stock.

**Table S3.**
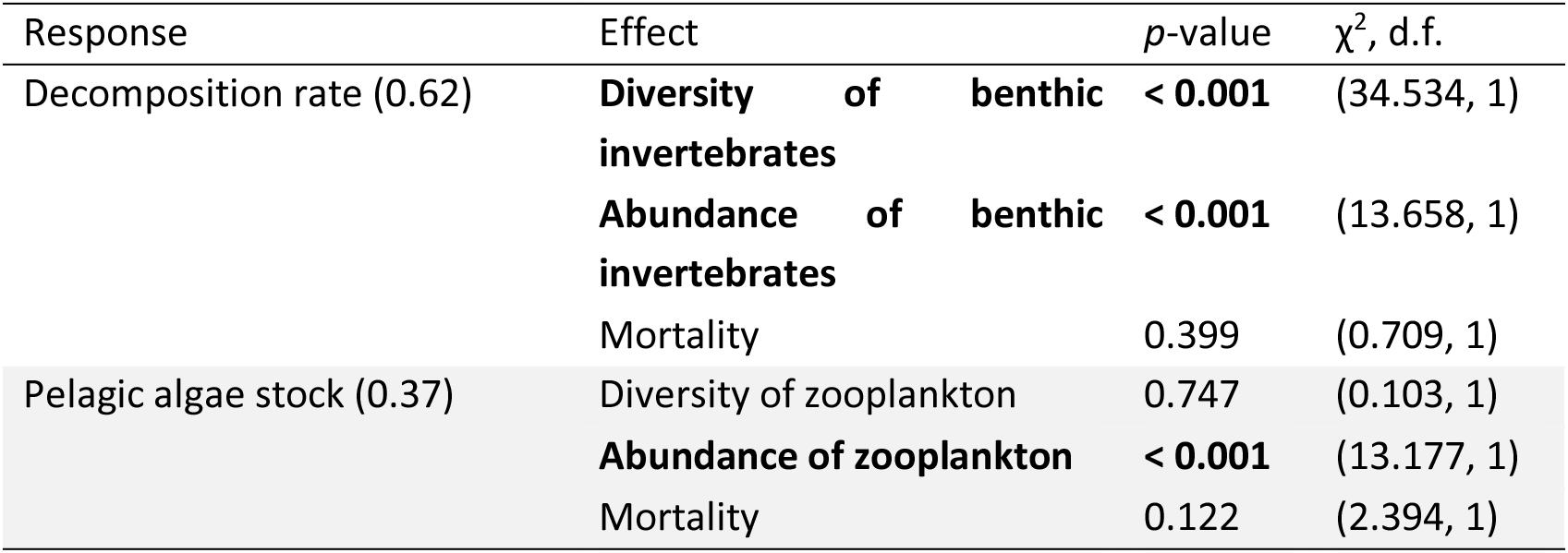
Linkages between community and ecosystem variables. Significant *p*-values are displayed in bold, R^2^ are shown into brackets.

## Notes

### Competing Interest Statement

The authors have declared no competing interest.

